# Phenotypic trait variation in a long-term, multisite common garden of Scots pine in Scotland

**DOI:** 10.1101/2022.06.08.495111

**Authors:** Joan Beaton, Annika Perry, Joan Cottrell, Glenn Iason, Jenni Stockan, Stephen Cavers

**Author notes:** corresponding author: Annika Perry.

## Abstract

Multisite common garden experiments, exposing common pools of genetic diversity to a range of environments, allow quantification of plastic and genetic components of trait variation. For tree species, such studies must be long term as they typically only express mature traits after many years. As well as evaluating standing genetic diversity, these experiments provide an ongoing test of genetic variation against changing environmental conditions and form a vital resource for understanding how species respond to abiotic and biotic variation. Finally, quantitative assessments of phenotypic variation are essential to pair with rapidly accumulating genomic data to advance understanding of the genetic basis of trait variation, and its interaction with climatic change.

We describe a multisite, population-progeny, common garden experiment of the economically and ecologically important tree species, Scots pine, collected from across its native range in Scotland and grown in three contrasting environments. Phenotypic traits, including height, stem diameter and budburst were measured over 14 growing seasons from nursery to field site. The datasets presented have a wide range of applications.

## Background & Summary

The need for comprehensive empirical assessments of genetic variation in tree species has never been greater. There is great interest around the world in growing more trees^1^, for their carbon sequestration abilities in the race for ‘net zero’ carbon emissions, to arrest biodiversity loss and forest decline^2^, and to manage watersheds^3^, as well as for the products they provide. However, there is considerable uncertainty in the accuracy of forecasts of future climate and the responses of tree species to those changes^4^. Intraspecific genetic variation and phenotypic plasticity will play key roles in determining how resilient existing and new tree populations are to the challenges ahead^5^. We are in urgent need of robust empirical data to calibrate the relationships between genetic variation within species and the environmental variables that will define future climates^6^. In parallel, the genomic revolution has provided a dramatic increase in the accessibility and scale of molecular data, and trees can now be genotyped faster, more cheaply and in greater number than ever before. However, there is a limit to what these new data can tell us without objective evaluation of the associated phenotypes, for which common garden and reciprocal transplant approaches remain key experimental tools^7,8^. We aimed to link these genomic and common garden approaches to better understand the genetic basis of phenotypic variation in trees, and to improve forecasting of how tree species will respond to climate change. Here we describe the research platform that we established to conduct these urgently needed long-term studies.

Scots pine (*Pinus sylvestris* L.) is globally very widely distributed, occurring predominantly from eastern Europe and Scandinavia to eastern Siberia, but with substantial populations in Scotland, southern Europe, Turkey and the Caucasus^9^. In Scotland, the remnant native populations, known locally as the Caledonian pinewoods, are typically small and highly fragmented and are distributed across a highly heterogeneous landscape that varies from oceanic (mild, very wet) environments in the west to more continental (drier, colder) in the east (Fig.1). This steep gradient of variation on a short spatial scale bears comparison to environmental gradients over much wider spatial scales across Europe (see Fig. 4 in Metzger et al., 2005^10^. To further the conservation of the pinewoods, specific action plans have been developed, including management of seed movements through a series of seven seed zones (Fig. 2^11^), the context and motivation for which are covered by Salmela et al., 2010^12^.

**Figure 1.**
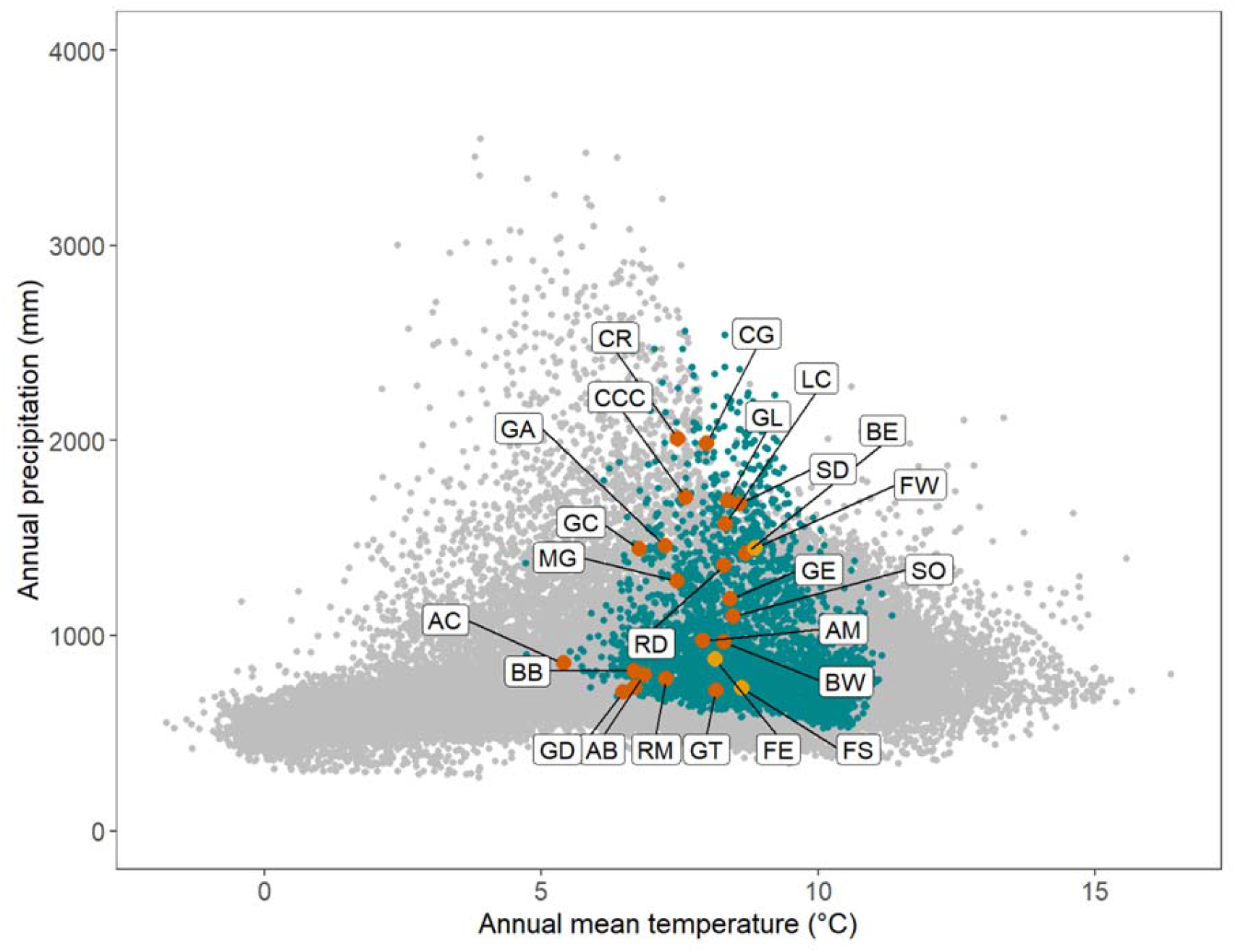
Ordination of 21 source populations in environmental space. Gray dots: global Scots pine distribution (distribution - EU-Forest database ^15^ and associated climate data (CHELSA database^16^, https://chelsa-climate.org/). Blue dots: Scots pine distribution in the United Kingdom. Orange dots: populations sampled for the experiment; letter codes match those in Table 1. Yellow dots: trial sites.

**Figure 2:**
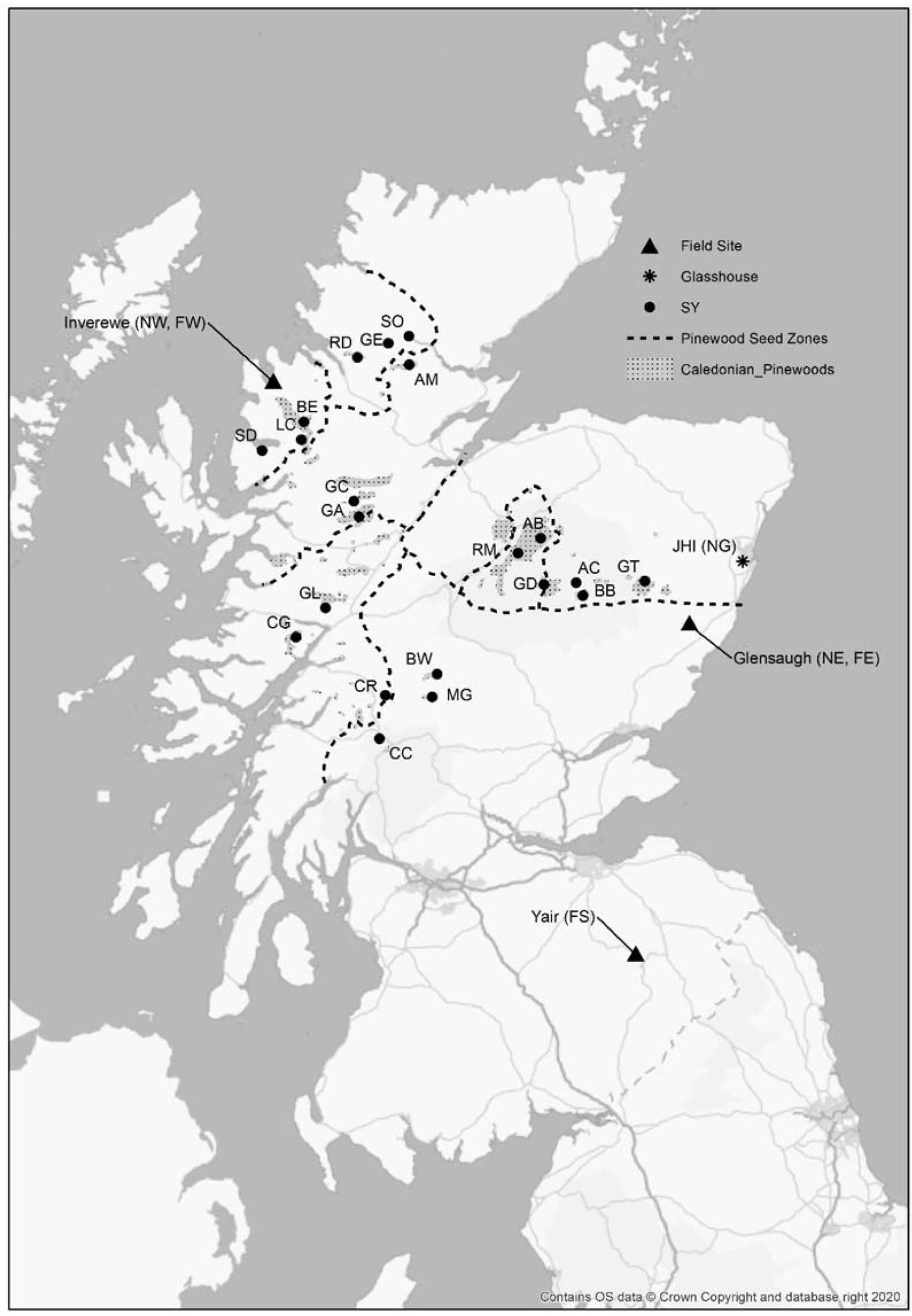
Location of the 21 source populations sampled for the experiment; letter codes match those in Table 1. Also shown are James Hutton Institute (JHI), location of eastern nursery (NE) and glasshouse (NG); three field sites - Inverewe (FW, also location of western nursery: NW), Glensaugh (FE) and Yair (FS).

A key knowledge gap remained in the early 21st century, namely the extent to which the current seed zone system for managing seed sourcing for the native populations of Scottish Scots pine reflects genuine genetic differences, and whether the zones conform to patterns of adaptation which are of relevance to ecological conservation and forestry when assessed using traits such as growth, mortality, phenology and resistance to pests and pathogens. To address this gap, as well as to evaluate the likely responses of indigenous Scots pine to ongoing climate change, an extensive, long-term assessment of genetic variation in Scots pine under contrasting environmental conditions, was founded by the Macaulay Institute (now the James Hutton Institute). A collaboration with the UK Centre for Ecology & Hydrology and Forest Research was subsequently created to develop the studies of adaptation to climate and disease resistance, to apply and advance emerging methods for assessing genetic variation, and to secure the continuation of the study. At the outset, an explicit objective was to sample genetic diversity widely without favouring any particular form or trait, and to undertake a wide ranging evaluation of traits covering different life history dimensions as far as practically possible. The common environment study was also intended to provide a long term experimental platform to facilitate future studies of the basis for variation in the extended phenotype of Scots pine including associated assemblages of organisms and community function.

**Table 1:** Locations and basic environmental data for the populations sampled for seed to establish the trial. See the maternal traits dataset (doi.org/10.5285/ac687a66-135e-4c65-8bf6-c5a3be9fd9aa) for precise data for each mother tree sampled.

| Population | Lat. (N) | Long. (W) | Alt. (m) | Area (ha) | GSL | GDD | FMT (°C) | JMT (°C) |
| --- | --- | --- | --- | --- | --- | --- | --- | --- |
| Abernethy (AB) | 57.21 | 3.61 | 311-370 | 2452 | 211 | 990 | 1.1 | 12.7 |
| Allt Cul (AC) | 57.04 | 3.35 | 435-512 | 13 | 145 | 513 | -1.0 | 10.4 |
| Amat (AM) | 57.87 | 4.60 | 39-201 | 181 | 214 | 892 | 1.2 | 12.3 |
| Ballochbuie (BB) | 56.98 | 3.30 | 421-531 | 775 | 116 | 446 | -1.7 | 9.5 |
| Beinn Eighe BE) | 57.63 | 5.40 | 17-91 | 182 | 283 | 1329 | 3.7 | 14.2 |
| Black Wood of Rannoch (BW) | 56.68 | 4.37 | 250-321 | 1011 | 254 | 1138 | 2.1 | 13.5 |
| Coille Coire Chuilc (CCC) | 56.42 | 4.71 | 222-311 | 67 | 226 | 928 | 1.6 | 12.3 |
| Conaglen (CG) | 56.79 | 5.33 | 89-193 | 189 | 246 | 887 | 2.2 | 11.7 |
| Crannach (CR) | 56.58 | 4.68 | 258-338 | 70 | 231 | 1019 | 1.8 | 12.6 |
| Glen Affric (GA) | 57.26 | 4.92 | 205-293 | 1532 | 210 | 769 | 0.9 | 11.6 |
| Glen Cannich (GC) | 57.35 | 4.95 | 182-381 | 301 | 212 | 778 | 1.0 | 11.7 |
| Glen Derry (GD) | 57.03 | 3.58 | 426-493 | 235 | 168 | 593 | -0.5 | 11.3 |
| Glen Einig (GE) | 57.96 | 4.76 | 45-92 | 27 | 242 | 1089 | 2.2 | 13.2 |
| Glen Loy (GL) | 56.91 | 5.13 | 136-219 | 74 | 191 | 541 | 0.5 | 9.8 |
| Glen Tanar (GT) | 57.02 | 2.86 | 289-422 | 1564 | 235 | 1105 | 2.2 | 13.6 |
| Loch Clair (LC) | 57.56 | 5.36 | 98-166 | 126 | 277 | 1253 | 3.4 | 13.7 |
| Meggernie (MG) | 56.58 | 4.35 | 254-385 | 277 | 223 | 916 | 1.1 | 12.0 |
| Rhidorroch (RD) | 57.89 | 4.98 | 138-220 | 103 | 221 | 840 | 1.5 | 11.6 |
| Rothiemurchus (RM) | 57.15 | 3.77 | 295-329 | 1744 | 224 | 1087 | 1.4 | 13.1 |
| Shieldaig (SD) | 57.50 | 5.63 | 44-132 | 103 | 273 | 1093 | 3.2 | 12.8 |
| Strath Oykel (SO) | 57.98 | 4.61 | 35-160 | 14 | 257 | 1276 | 2.7 | 14.0 |

Alt. - altitudinal range sampled within each population, Area - core pinewood area<sup>13</sup>. Average (1961-2000) climate variables from UK Met Office<sup>14</sup>; GSL - growing season length (days), GDD - growing degree days (day degrees), FMT - February and JMT - July mean temperatures (°C).

The following describes the origins, design and initial measurements of a multi-site experiment in Scots pine, including protocols adopted during both the initial nursery phase and the final field experiment.

## Methods

### Seed sampling and germination

Seed from ten trees from each of 21 native Scottish Scots pine populations (Table 1) were collected in March 2007 and germinated at the James Hutton Institute, Aberdeen (latitude 57.133214, longitude -2.158764) in June 2007. Populations were chosen to represent the species’ native range in Scotland and to include three populations from each of the seven seed zones (Fig. 2). There was no selection of seed-trees on the basis of any traits except for the possession of cones on the date of sampling. Ten seed trees were sampled from each population according to a spatial protocol designed to cover a circle of approximately 1 km in diameter located around a central tree. The sampling strategy identified nine points each in a pre-determined random direction from the central point, whilst stratifying the number sampled with increasing distance from the central point in the ratio 1: 3: 5. This strategy avoids over-sampling the areas close to the centre point. For smaller fragments of woodland, or where only a few trees with cones were present, then the directions of the sampled trees from the central tree were maintained to give a wide coverage of the woodland area, but the distances between trees varied but were never closer than 50 m. To break dormancy, seeds were soaked for 24 hours on the benchtop at room temperature, after which they were stored in wet paper towels and refrigerated in darkness at 3-5 °C for approximately 4 weeks. Seeds were kept moist and transferred to room temperature until germination began (approx. 5-7 days), then transplanted to 8 cm x 8 cm x 9 cm, 0.4 L pots filled with Levington’s C2a compost and 1.5g of Osmocote Exact 16-18 months slow release fertiliser and kept in an unheated glasshouse. The compost was covered with a layer of grit to reduce moss and liverwort growth. Seedlings from the same mother tree are described as a family and are assumed to be half-siblings.

### Experimental design: nurseries

The full collection consisted of 210 families (10 families from each of 21 populations) each consisting of 24 half sibling progeny (total 5,040 individuals); needle tissue was sampled from each seedling and preserved for long term storage, one needle on silica gel, 2-5 needles at - 20 °C. After transfer into pots, 8 seedlings per family were moved to one of three nurseries (total 1,680 seedlings per nursery): outdoors at Inverewe Gardens in western Scotland (nursery in the west of Scotland: coded NW, latitude 57.775714, longitude -5.597181, Fig. 2); outdoors in a fruit cage (to minimise browsing) at the James Hutton Institute in Aberdeen (nursery in the east of Scotland: NE); in an unheated glasshouse at the James Hutton Institute in Aberdeen (nursery in a glasshouse: NG). Trees were arranged in 40 randomised trays (blocks) in each nursery. Each block contained two trees per population (total 42 trees). Watering was automatic in NG, and manually as required for NE and NW. No artificial light was used in any of the nurseries. In May 2010 the seedlings from NG were moved outdoors to Glensaugh in Aberdeenshire (latitude 56.893567, longitude -2.535736). In 2010 all plants were repotted into 19 cm (3 L) pots containing Levingtons CNSE Ericaceous compost with added Osmocote STD 16-18 month slow release fertilizer.

### Experimental design: field sites

In 2012 the trees were transplanted to one of three field sites: Yair in the Scottish Borders (field site in the south of Scotland: FS, latitude 55.603625, longitude -2.893025); Glensaugh (field site in the east of Scotland: FE); and Inverewe (field site in the west of Scotland: FW). All trees transplanted to FS were raised in the NG and all but four of the trees transplanted to FE were raised locally in the NE (the remainder were grown in NG). In contrast, following mortality and ‘beating up’ (filling gaps where saplings had died), the FW experiment ultimately contained cohorts of trees raised in each of the three nurseries as follows: 290 grown locally in the NW; 132 were grown in the NG; and 82 were grown in the NE.

#### Site histories

The Yair site (FS) had previously been used for growing Noble fir (*Abies procera*) for Christmas trees and Lodgepole pine (*Pinus contorta*), a section of the former were felled and chipped to create a clear area prior to planting. The planting site is also adjacent to a large block of commercial Sitka spruce (*Picea sitchensis*) forestry, and the Glenkinnon Burn Site of Special Scientific Interest (SSSI naturescot site code 736; EU site code 135445), an area of mixed broadleaf woodland. Prior to planting, major areas of tall weeds were strimmed. The site was protected by a deer fence. The experiment was planted 8-11 October 2012. The Glensaugh site (FE) is in Forestry Compartment 3 of the Glensaugh Research Station, adjacent to Cleek Loch. It is thought to have been cleared of Scots pine and Larch (*Larix decidua*) around 1917, after which it reverted to rough grazing. An attempt to reseed part of the site in the 1980s was unsuccessful and it quickly reverted to rough grazing for a second time. The whole site (within which the experimental area is embedded) was deer fenced and re-planted under the Scottish Rural Development Programme (SRDP) in 2012. The experimental plot was planted up 7-9 March 2012. The Inverewe site (FW) had previously been a Sitka spruce and Lodgepole pine plantation (50:50 mix) that had been clear-felled in 2010 following substantial windthrow. The experimental site was deer fenced in early 2012, and the experiment was planted 12-16 March 2012, followed by beating up on 27-28 March 2013 and 22-24 October 2013. There had been minimal preparation of the site in line with current practice for restocking sites. The experimental site is included in the Inverewe Forest Plan, which included deer fencing of a larger area (2014) around the experimental site. Planting of this area was be completed in early 2015, funded by NTS (National Trust for Scotland), although natural regeneration is also taking place.

At each site, trees were planted in randomised blocks at 3 m x 3 m spacing. There are four randomised blocks in both FS and FE and three in FW. A guard row of Scots pine trees was planted around the periphery of the blocks. Each block comprised one individual from each of eight (of the 10 sampled) families per 21 populations (168 trees). Although most families (N = 159) were represented at each of the three sites, families with insufficient trees (N = 9) were replaced in one site (FS) with a different family from the same population. Each experimental site was designed with redundancy such that, if thinning becomes necessary as the trees mature, then the systematic removal of trees (i.e. trees 1,3,5,7, etc of row 1, and 2,4,6,8, etc of row 2, 1,3,5,7,etc of row 3) will maintain a balanced design of the experiment, with sufficient family and population representation to provide an ongoing experiment with full geographic coverage.

The field sites generally experience different climates, with FW typically warmer and wetter and with more growing degree days per year and a much longer growing season than both FE and FS (Table 2). The coldest site with the shortest growing season is generally FE.

**Table 2.**
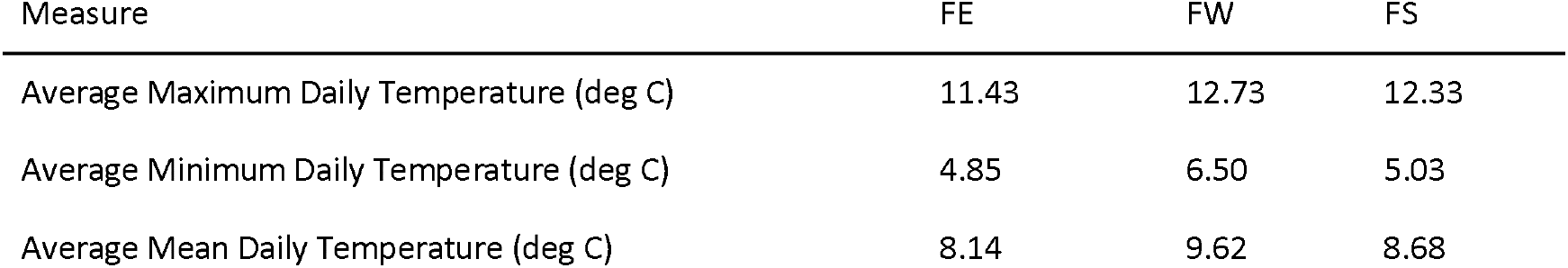
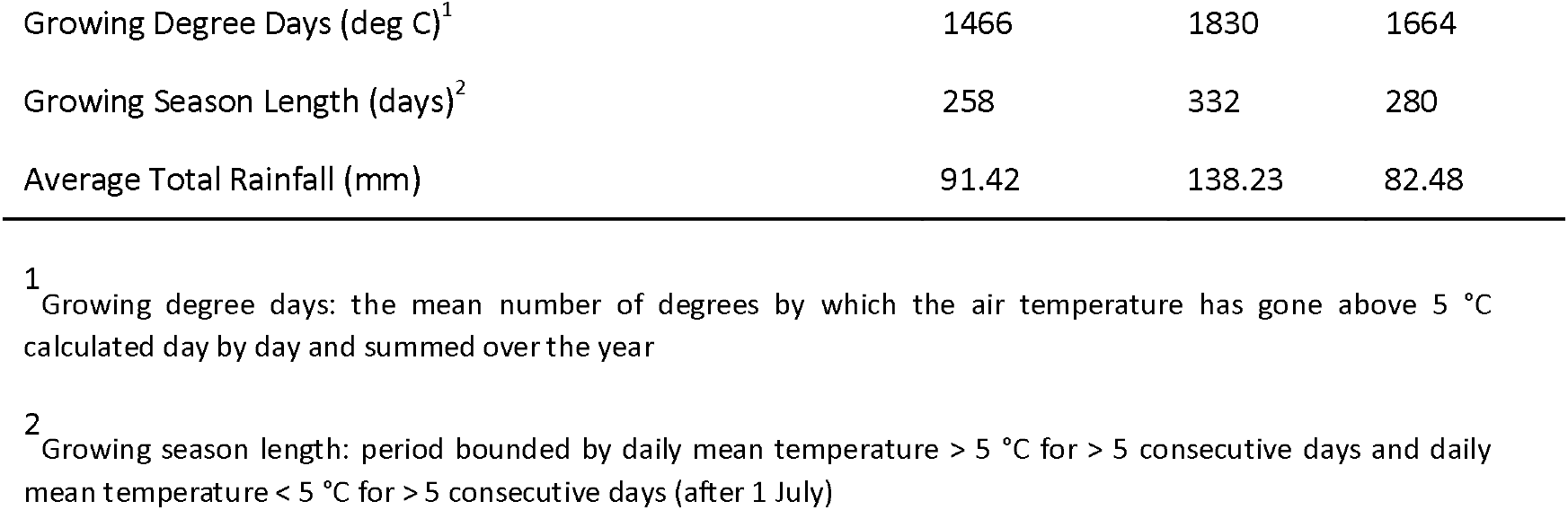
Average climatic variables at Glensaugh (FE), Inverewe (FW) and Yair (FS) from planting in 2012 until 2020. Climatic variables are derived from data provided by the Met Office (daily mean, minimum and maximum temperatures and monthly rainfall).

### Phenotype assessments

#### Maternal traits

Following seed collection, a range of traits were measured in the mother trees in order to control for maternal effects in subsequent measurements of their progeny (Table 3). For each mother tree, measurements of height and diameter at breast height (DBH) were taken, and ten cones were collected and assessed in detail. Cone width and length were measured prior to drying the cones (when they were still closed). Cone weight was measured post-drying. Seed removed from each cone was assessed for total weight and for the count and percentage of seeds which were classed as viable (viable seed were those which had both a wing and an obvious seed).

**Table 3:** Traits assessed in mother trees, cones, seeds, nursery seedlings and field trials. Within the datasets, traits are recorded in a single column for each year using the format Code-Year (e.g. absolute height in 2008 = HA08) except for the maternal traits datasets which were all measured in 2007. Where multiple measurements are made in a single year (i.e. for canopy width and needle length) the suffix “_X” is added to the column header, where X is the measurement number (e.g. canopy width measured in 2007, second measurement = CW07_2). Where multiple stages are recorded in a single year (i.e. for budburst timing) the suffix “_Y” is added to the column header, where Y is the budburst stage.

| Trait | Code | Unit | Dataset | Year(s) |
| --- | --- | --- | --- | --- |
| Cone length | Le | mm | Maternal | 2007 |
| Cone weight | We | G | Maternal | 2007 |
| Cone width | Wi | mm | Maternal | 2007 |
| Height: absolute | HA | M<br>mm<br>mm | Maternal<br>Nursery<br>Field | 2007<br>2007-2011<sup>A</sup><br>2013-2020<sup>B</sup> |
| Height: increment | HI | mm<br>mm | Nursery<br>Field | 2008-2011<sup>A</sup> |
| Viable seed number | SN | Count | Maternal | 2007 |
| Viable seed weight | SW | G | Maternal | 2007 |
| Seed viability | SP | % | Maternal | 2007 |
| Stem diameter: absolute<sup>C</sup> | DA | M<br>mm<br>mm | Maternal<br>Nursery<br>Field | 2007<br>2008-2011<sup>A</sup><br>2013-2020<sup>D</sup> |
| Stem diameter: increment | DI | mm<br>mm | Nursery<br>Field | 2009-2011<sup>A</sup> |
| Canopy width | CW | mm | Nursery | 2008 |
| Growth cessation | GC | Days since 1 Sep 2008 | Nursery | 2008 |
| Needle length | NL | mm | Nursery | 2007-2009 |
| Number of buds | Bu | Count | Nursery | 2008 |
| Budburst duration | BD | Days | Field | 2015-2019 |
| Budburst timing | BB<br>BT | Days since 31 March | Nursery<br>Field | 2008<br>2015-2019 |

#### Nursery traits

Seedling phenotype assessments were performed annually from 2007-2010 for three different trait types: phenology (budburst and growth cessation); form (total number of buds, needle length); cumulative growth (stem diameter and height, canopy width). Measurements of tree form and cumulative growth traits were taken after the end of each growing season. Phenology was assessed weekly during the spring and autumn of 2008 for budburst and growth cessation, respectively. Budburst was defined as the number of days from 31 March 2008 to the time when newly emerged green needles were observed. Growth cessation was defined as the number of days from 1 September 2008 to the time when no further growth was observed. Canopy width (widest point) was measured at two perpendicular points in the horizontal plane. Needle length was measured for three needles per tree. Mortality was recorded each year from 2007 to 2010.

#### Field traits

Tree height was measured in the field in the winter after each growing season from 2013 at FE and FW, and from 2014 to 2020 at all sites. Height was taken as the vertical measurement in cm from top bud straight to the ground. Basal stem diameter was measured at the end of the growing season for trees growing at FE and FW from 2014 to 2020 and for FS in 2020.

Phenology assessments were performed in spring at each site from 2015 to 2019. Seven distinct stages of budburst were defined (Supplementary Table 1) although only stages 4 to 6 are considered for analysis due to high proportions of missing data for the early and late stages. Each tree was assessed for budburst stage at weekly intervals from early spring until budburst was complete, annually from 2015 until 2019. In order to allow comparisons within and among sites and years, the date at which each stage of budburst occurred was considered relative to 31 March of that year. For example, 25 May 2019 is recorded as 55 days since 31 March 2019. The duration of budburst (time taken to reach stage 6 from stage 4) was also estimated.

When trees progressed through budburst stages rapidly, skipping a stage between assessments, a mean value was taken from the two assessment dates. For example, if a tree was at stage 4 on day 55 and was recorded as stage 6 at the next assessment on day 62, it is assumed to have reached stage 5 at day 58.5.

## Data Records

Data are deposited with the Environmental Information Data Centre (http://eidc.ac.uk)

DOI for maternal traits datasets: doi.org/10.5285/ac687a66-135e-4c65-8bf6-c5a3be9fd9aa

DOI for nursery traits dataset: doi.org/10.5285/29ced467-8e03-4132-83b9-dc2aa50537cd

DOI for field traits dataset: doi.org/10.5285/f463bc5c-bb79-4967-a8dc-f662f57f7020

In *all* datasets, the first two columns are:

1: Population code (code for forest of origin, 21 total)

2: Family (unique mother tree code: progeny described in nursery traits dataset and field trait dataset from the same family are putative half-siblings)

There are two maternal traits datasets: one for traits relating directly to the mother tree (MotherTraits.txt) in which each row represents one tree, and a second for traits relating to cones and seed collected from each mother tree in which each row represents one cone collected from each tree. Columns are defined as follows:

*Maternal dataset*: MotherTraits.txt

3: Population (name of forest of origin, 21 total)

4: Seed zone

5-13. Location reference and immediate environment for each tree: Latitude [decimal]; Longitude [decimal]; Aspect; Slope; Altitude [m]; Regeneration [1-4]; Peat depth [cm]; soil moisture [1-5]; mean distance to nearest three trees [M]

14-15. Mother tree traits (absolute height [M]; diameter at breast height [M])

*Maternal dataset*: ConeSeedTraits.txt

3. Cone number (1-10)

4-6. Cone traits (Width [mm], Length [mm], Weight [g])

7-9. Seed traits (Viable seeds [count], Percentage viable seeds (%), Viable seeds weight [g])

*Nursery traits dataset*:

3. Seedling id

4. Nursery site [NE; NW; NG]

5-6. Nursery block number: from 2007 to 2010 [1-40]; from 2010 to 2012 [1-98]

7. Field code if transplanted [4 digit code; not transplanted = NA]

8-40. Nursery traits (see Table 3 for list of traits and format of column header)

41-44. Status [Alive; Dead] for each year 2007-2010 (inclusive)

*Field traits dataset*:

3. Field site [FE; FS; FW]

4. Field code [4 digit code]

5. Block number [A; B; C; D]

6-56. Field traits (see Table 3 for list of traits and format of column header)

## Technical Validation

Measurements repeated annually are performed at the same time of year to ensure consistency of the method, e.g. height is measured during winter to avoid the possibility of active growth occurring after the trees are measured. Data were checked after each survey and inconsistent values (i.e. where height was less than the previous year) were re-measured. Where height increment was found to be less than 0 mm (due to an error in measurement or an effective loss in height due to damage) values were removed and classed as missing (NA). Annual stem diameter increment was estimated as the increase in stem diameter from the end of one growing season to the end of the next. Where stem diameter increment was estimated between 0 and -2 mm, the error was assumed to be caused by differences in orientation of measurement between years and increment values were adjusted to 0. Where stem diameter increment was found to be less than -2 mm, the increment and most recent stem diameter measurements were both classed as missing (NA).

We used boxplots to visualize data range and data distribution for each trait in each year over all nursery and field sites and all populations (Fig. 3). The use of outliers in subsequent analyses should be treated with caution.

**Figure 3.**
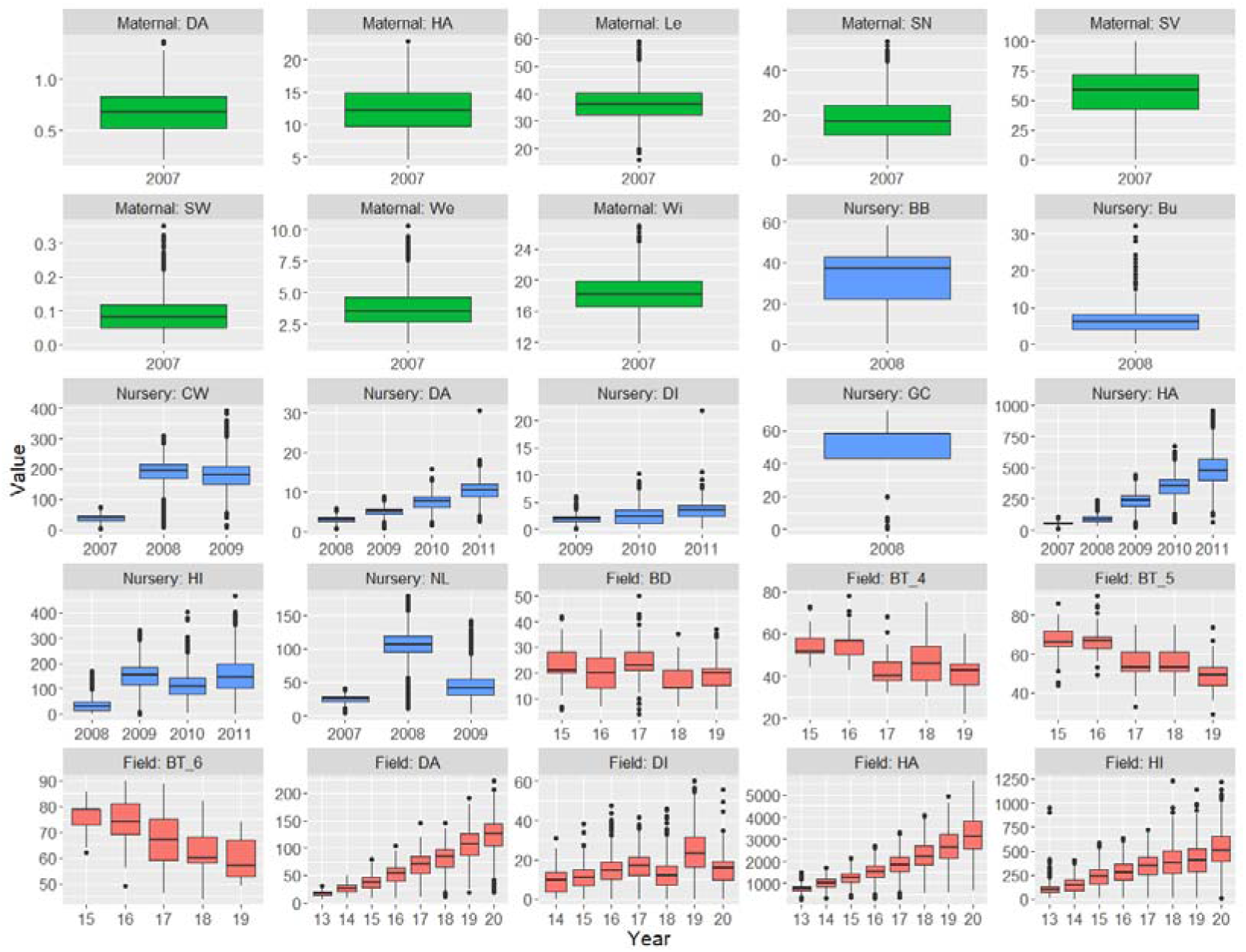
Box and whisker plots of trait values for each trait in each year over all nursery/field sites and all populations. Trait value units are listed in Table 3. Year for field dataset traits are abbreviated (e.g. 2015 = ‘15’). Solid black lines indicate the median trait value. The bottom and top of boxes indicate the first and third quartile. The upper and lower whiskers extend to the highest and lowest values within 1.5 times the interquartile range. Individual points indicate outliers. Traits derived from the maternal datasets are green, those from the nursery dataset are blue and those from the field dataset are red.

## Acknowledgements

The James Hutton Institute contribution to this work was funded by Scottish Government’s Rural & Environment Science & Analytical Services Division, through their Strategic Research Programmes (2006-2011, 2011-2016 and 2016–2022). We thank Stewart Mackie and staff at Forest and Land Scotland (Selkirk Office); the Commission /National Trust for Scotland, and Donald Barrie (Glensaugh) for their help in locating, establishing and securing access to sites. A large number of people have contributed to the design, establishment, maintenance and monitoring of this experiment, these include Dave Sim, Ben Moore, Jack Lennon, Luis Miguel Quinzo Ortega, Patrick McKenzie, Douglas Iason, Sheila Reid, Diana Menzel, Francisco Pou-Bolibar, Sarah Williams, Vivian Bisset, Debbie Fielding, Richard Hewison, Isla Morton, Alison Hester, Rutger Scholtens, Alan Gray, Kevin Donnelly, Nikoleta Ntiantiasi, Michelle Biermann, Juan Pablo Lobo-Guerrero. We also thank Richard Ennos, Pete Hollingsworth and Julianne O’Reilly-Wapstra for discussions that contributed to the establishment of the project.

## Author contributions

GI originated and initiated the project, including design of the seed collection and nursery experiment, and design of the field experiment, along with SC (and assistance of Betty Duff, BioSS). JB performed seed collections, nursery and transplanting work, and collated data and managed the nursery experiment and the FE and FW field sites. SC, AP, JC planted and managed the FS site. All authors contributed to data collection and maintenance of the field experiments.

## Competing interests

The authors declare no conflict of interest

**Supplementary Table 1:**
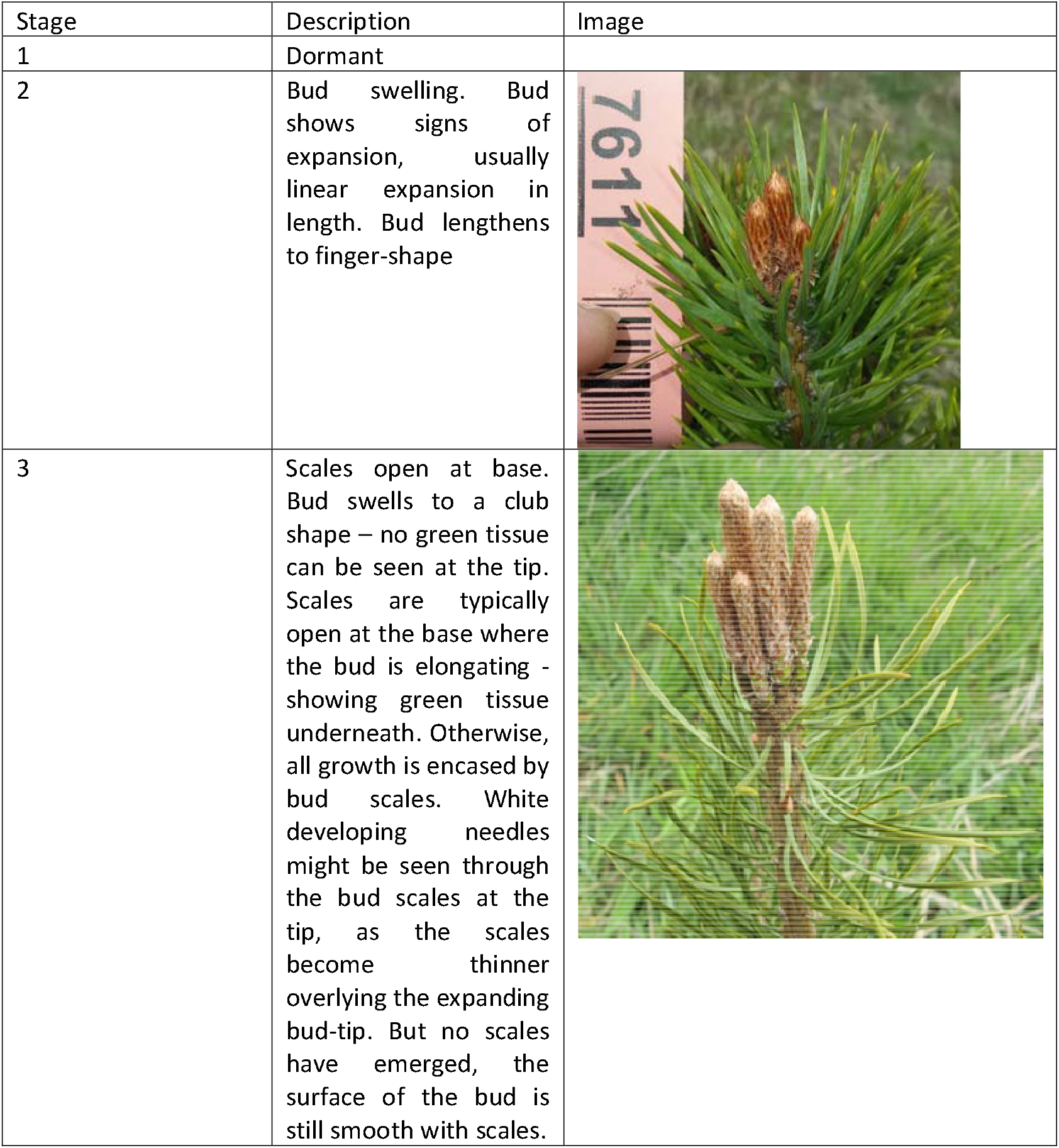
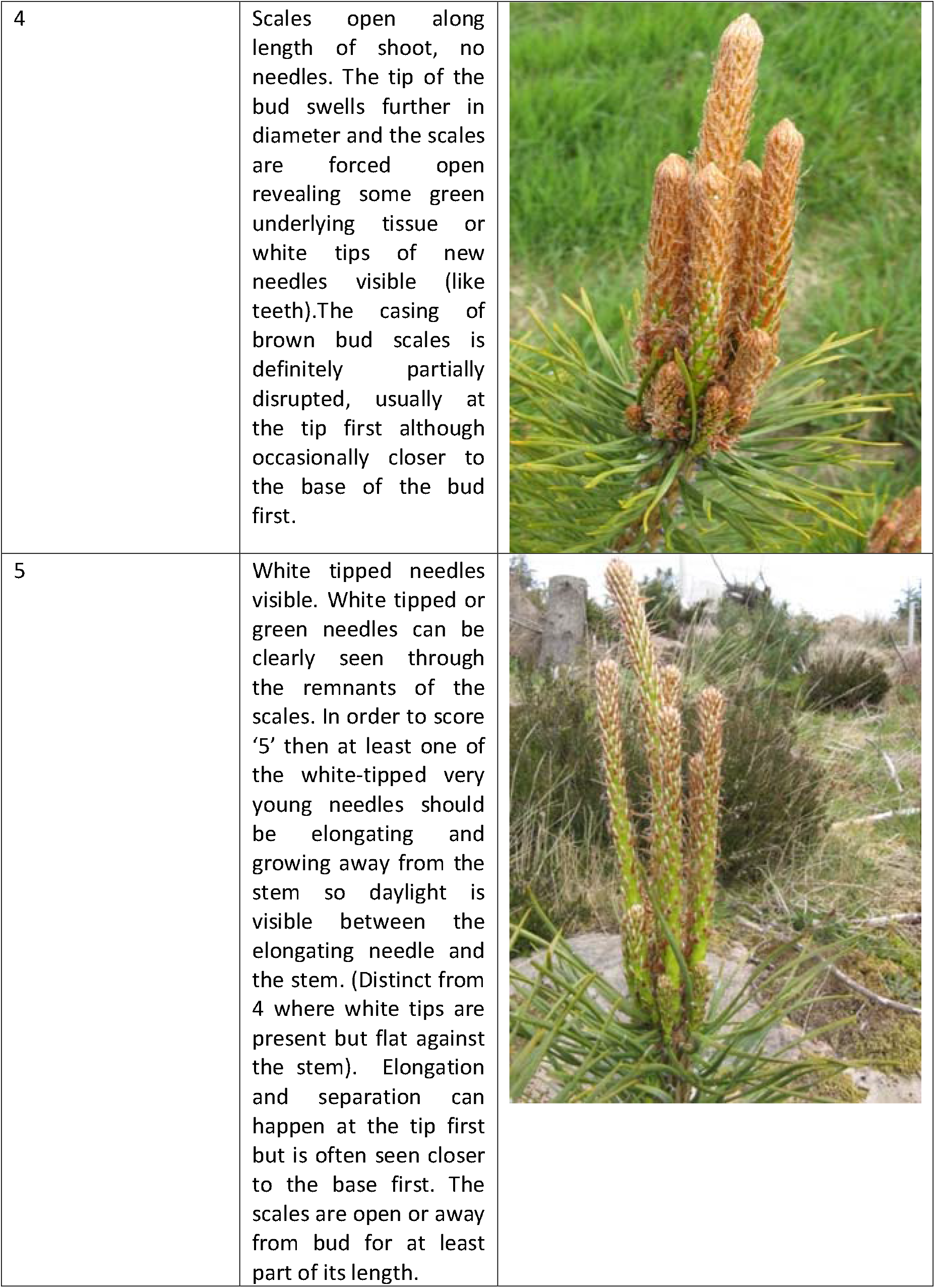
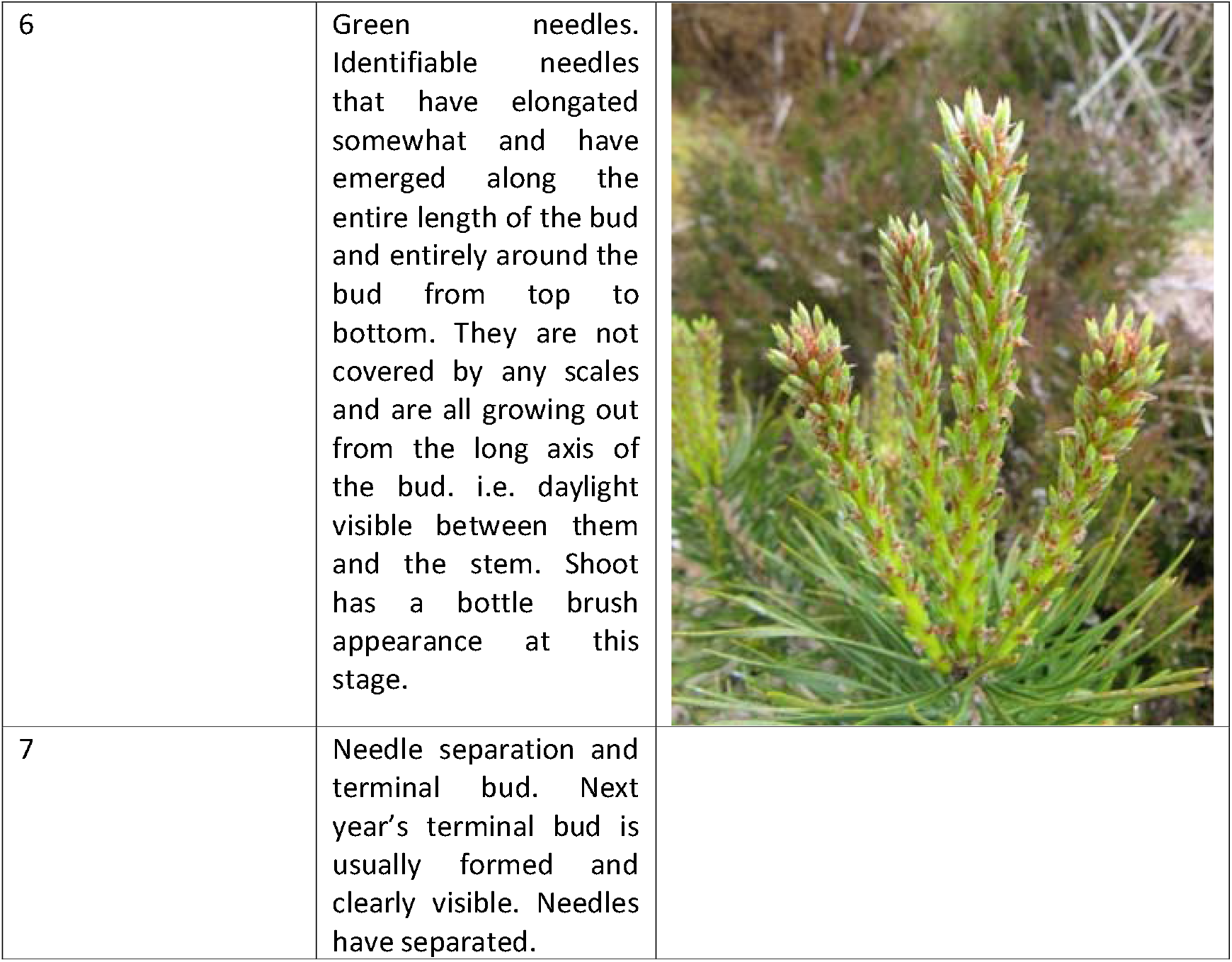
Phenological stages of bud burst assessed in field trials.

| Stage | Description | Image |
| --- | --- | --- |
| 1 | Dormant |  |
| 2 | Bud swelling. Bud shows signs of expansion, usually linear expansion in length. Bud lengthens to finger-shape |  |
| 3 | Scales open at base. Bud swells to a club shape – no green tissue can be seen at the tip. Scales are typically open at the base where the bud is elongating - showing green tissue underneath. Otherwise, all growth is encased by bud scales. White developing needles might be seen through the bud scales at the tip, as the scales become thinner overlying the expanding bud-tip. But no scales have emerged, the surface of the bud is still smooth with scales. |  |
| 4 | Scales open along length of shoot, no needles. The tip of the bud swells further in diameter and the scales are forced open revealing some green underlying tissue or white tips of new needles visible (like teeth). The casing of brown bud scales is definitely partially disrupted, usually at the tip first although occasionally closer to the base of the bud first. |  |
| 5 | White tipped needles visible. White tipped or green needles can be clearly seen through the remnants of the scales. In order to score '5' then at least one of the white-tipped very young needles should be elongating and growing away from the stem so daylight is visible between the elongating needle and the stem. (Distinct from 4 where white tips are present but flat against the stem). Elongation and separation can happen at the tip first but is often seen closer to the base first. The scales are open or away from bud for at least part of its length. |  |
| 6 | Green needles. Identifiable needles that have elongated somewhat and have emerged along the entire length of the bud and entirely around the bud from top to bottom. They are not covered by any scales and are all growing out from the long axis of the bud. i.e. daylight visible between them and the stem. Shoot has a bottle brush appearance at this stage. |  |
| 7 | Needle separation and terminal bud. Next year's terminal bud is usually formed and clearly visible. Needles have separated. |  |

